# Identifying Neural Signatures Mediating Behavioral Symptoms and Psychosis Onset: High-Dimensional Whole Brain Functional Mediation Analysis

**DOI:** 10.1101/2020.04.15.043034

**Authors:** Oliver Y. Chén, Hengyi Cao, Huy Phan, Guy Nagels, Jenna M. Reinen, Jiangtao Gou, Tianchen Qian, Junrui Di, John Prince, Tyrone D. Cannon, Maarten de Vos

## Abstract

Along the pathway from behavioral symptoms to the development of psychotic disorders sits the multivariate mediating brain. The functional organization and structural topography of large-scale neural mediators among patients with brain disorders, however, are not well understood. Here, we design a high-dimensional brain-wide functional mediation framework to investigate brain regions that intermediate between baseline behavioral symptoms and future conversion to full psychosis among individuals at clinical high risk (CHR). Using resting-state functional magnetic resonance imaging (fMRI) data from 263 CHR subjects, we extract an ***α*** brain atlas and a ***β*** brain atlas: the former underlines brain areas associated with prodromal symptoms and the latter highlights brain areas associated with disease onset. In parallel, we identify the *P* mediators and the *N* mediators that respectively facilitate or protect against developing brain disorders among subjects with more severe behavioral symptoms and quantify the effect of each neural mediator on disease development. Taken together, the ***α***-***β*** atlases and the *P*-*N* mediators paint a brain-wide picture of neural markers that are potentially regulating behavioral symptoms and the development of psychotic disorders and highlight a statistical framework that is useful to uncover large-scale intermediating variables in a regulatory biological organization.

## INTRODUCTION

How does the human brain intermediate between behavioral symptoms and the development of brain diseases? Which brain areas are involved in this process? Can we chart these areas’ functional characteristics and structural organization?

Researchers studying brain diseases often observe that the patterns of the brain are on the one hand associated with behavioral symptoms, and on the other hand linked to disease status. Conventionally, the former is called an independent variable, the latter a dependent variable (or an outcome), and the brain patterns interposed in-between a mediator. A central problem in neural mediation analysis is to identify which brain regions are positioned along the causal pathway between behavioral symptoms and disease status. Equally important is to quantify the effect of each identified brain area on developing the disease and to determine its relative prominence in the causal hierarchy.

Disorganization symptoms, such as bizarre thoughts and behaviors, are considered to be associated with conversion to psychosis among individuals at clinical high risk (CHR); empirical studies have shown a significantly higher hazard ratio for psychosis onset in CHR subjects with higher disorganization symptoms at baseline^1–3^. Yet, as properties associated with a mental disorder, the disorganization symptoms and disease development are reflected by the patterns of the brain. By probing into the neural basis of human behavior and disease development, mediation analysis can help us to understand the functional organization and structural distribution of the brain patterns that regulate behavioral symptoms and disease development. But it can only do so by first charting the neural pathways that make brain mediation possible.

A beginning in this direction can be made by identifying and isolating neural mediators that are interposed between behavioral symptoms and disease development (see **Figure** 1). Neural mediators in psychosis studies are high-dimensional (involving patterns from hundreds of thousands of areas distributed across the brain), their patterns functional (they may consist of differentiable functional neural signals), and their corresponding outcomes binary (*e.g*. whether one has a full-blown psychotic disorder or not). One must therefore confront several key challenges to uncover high-dimensional functional neural mediators. First, although existing mediation models have made the search for mediators fruitful, they are not suitable for studying high-dimensional mediation analysis with binary disease outcomes. For example, existing multi-level mediation models assume that the outcomes are continuously distributed^4–7^; mediation frameworks concerning binary outcomes are at present restricted to small-scale mediators^4–6,9^; high-dimensional mediation models whose outcomes are not normally distributed do not have a closed form solution (therefore it is difficult to estimate parameters analytically)^see 4^. Second, although functional mediation analysis^8^ has advanced considerably knowledge about the functional signal organization of the brain in relation to independent and outcome variables, it remains unclear whether it is suitable for analyzing high-dimensional brain data, and if so, how the underlying data configuration, such as the sample size and noise level, would affect parameter estimation. In parallel, its efficacy needs to be evaluated for brain disease studies. Third, the underlying functional bases of the brain patterns are largely unknown and could be orthogonal or nonorthogonal; additionally, the data can be contaminated by noise. Whether and how the functional architecture of the neural bases, their orthogonality, and noise level, would affect mediation analysis is an as-of-yet less-well-charted area. If not properly treated, this set of circumstances could generate inconsistent methodological results and confusing interpretations. Additionally, if one manages to discover functional neural mediators, the discovery naturally raises the question of which mediators are facilitative, and which are protective against developing brain disorders among subjects with more severe behavioral symptoms.

**Figure 1.**
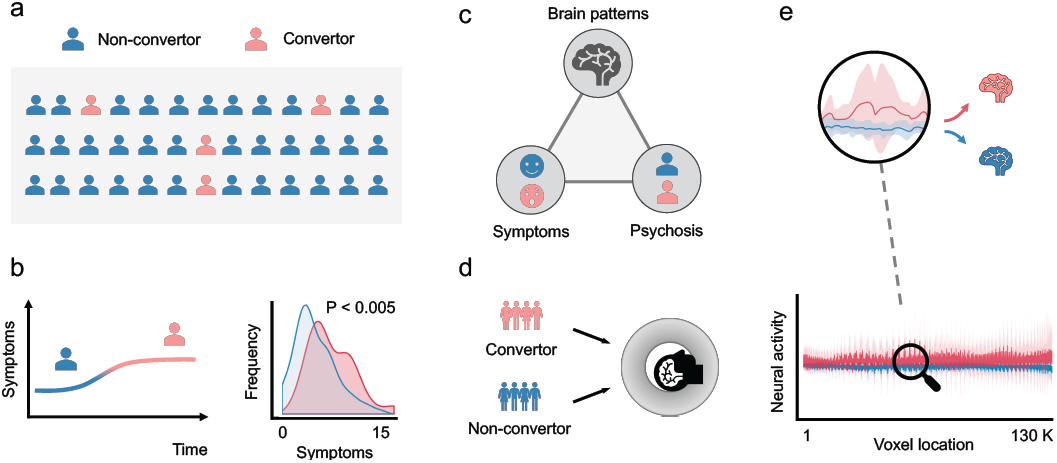
The study layout of the neural mediation analysis. (a) We considered a sample of 263 subjects recruited from eight study sites across the United States and Canada who met criteria for a prodromal risk syndrome at the point of recruitment and had been clinically followed up for two years as part of the NAPLS-2 project. During the follow-up period, 25 subjects developed a full-blown psychotic disorder (CHR convertors); 238 did not (CHR non-convertors). (b) The behavioral symptoms of convertors were significantly more severe than those of non-convertors. (c) The neural mediation analysis investigated which brain regions were intermediating between psychosis symptoms and disease status. Once neural mediators were identified, one could further quantify the mediation effect of each mediator to determine its relative prominence in the causal hierarchy. (d) Both convertors and non-convertors received an eyes-open resting-state functional magnetic resonance imaging (fMRI) scan at the point of recruitment. (e) The fMRI BOLD signals from both convertor and non-convertor samples were plotted along 130,992 brain areas. The red shade were BOLD signals stacked across the convertor group across the whole-brain space and the blue shade were from the non-convertor group.

To address these questions and challenges, we designed a high-dimensional functional mediation model. Through simulation studies and empirical data analysis, we demonstrate that the model is able to (a) analyze large-scale intermediating brain patterns (*e.g*., fMRI BOLD signals from hundreds of thousands of voxels); (b) distinguish distinctive functional patterns of the brain between disease groups in relation to behavior symptoms while accounting for covariates; (c) quantify each neural mediator’s effect on disease outcome; and (d) identify and separate brain areas that are facilitative or protective for developing brain disorders.

In the following, we begin with a brief overview of the mediation analytical frameworks concerning univariate and multivariate mediators. After discussing these basic concepts, we introduce the high-dimensional functional mediation framework. To demonstrate its utility, we perform both simulation and case studies. During the simulation study, we consider various experimental settings, including different levels of noise, sample sizes, and both orthogonal and non-orthogonal basis functions, to ensure that the proposed framework is suitable for studying high-dimensional functional mediation. During the case study, we uncover brain areas that regulate psychosis symptoms and disease status from whole-brain resting-state functional magnetic resonance imaging (rs-fMRI) data obtained from 263 subjects at clinical high risk (CHR) for psychosis.

### Univariate mediation analysis

Univariate mediation analysis considers a single mediator (*M*) (see **Figure 2** (a)). In other words, a variable *M* is a mediator if, after accounting for covariates ***Z***, the effect of an independent variable *X* on an outcome variable *Y* is at least partially carried through *M* ^10,11^. Examples of univariate mediators are pain catastrophizing, which mediates the clinical treatment (*X*) and disability status (*Y*)^12^, and intention, which mediates attitudes (*X*) and behavior (*Y*)^13^.

**Figure 2.**
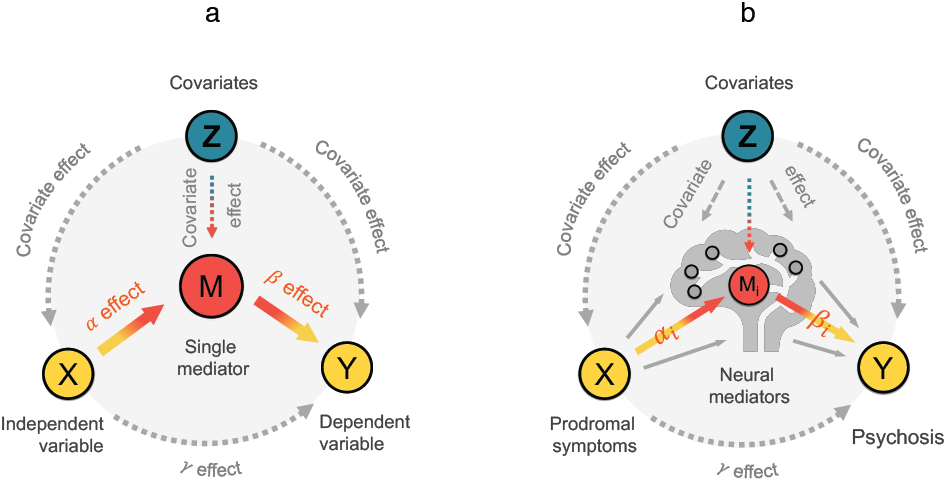
A schematic representation of multivariate mediation analysis. (a) Univariate mediation analysis. The circles indicate an independent variable, a univariate mediator, an outcome variable, and covariates. The arrows denote potential causal pathways. The letter *α* denotes the effect from the independent variable to the mediator, after accounting for the covariate effect. The letter *β* denotes the effect of the mediator on the outcome, after controlling the independent variable and covariates. The letter *γ* denotes the effect from the independent variable to outcome, after accounting for the covariate effect. (b) Multivariate neural mediation analysis. Each red circle within the brain represent a potential neural mediator. The arrows denote potential causal links. The letter *α*_*i*_ (1 ≤ i ≤ *V*) denotes the effect from the independent variable to the *i*^*th*^ neural mediator (represented by a red circle). The letter *β*_*i*_ denotes the effect of the neural mediator on the outcome, after controlling the independent variable and covariates. The letter *γ* indicates the direct effect from the independent variable to outcome, after accounting for the covariate effect.

The identification of a univariate mediator consists of two steps^10,14–17^ (see **Supporting Information** for a comparison between two common mediation analysis frameworks). The first step examines if the independent variable *X* has an effect on the mediator *M* after controlling for the covariates ***Z***, using the following conditional model:

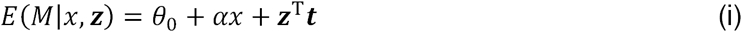

where *E* refers to the expectation operation; *θ*_0_, *α*, ***t*** are coefficients for the intercept, the observed independent variable *x*, and the observed covariates ***z***. If *α* is significantly different from zero, then the independent variable has an effect on the mediator.

The second step evaluates if the mediator *M* has an effect on the outcome *Y*, after controlling for the independent variable *X* and covariates ***Z***, using the following conditional model:

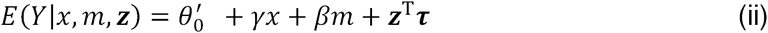

Where 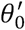, *γ, β* and **τ** are coefficients for the intercept, the observed independent variable *x*, the mediator *m*, and the observed covariates ***z***. The prime in 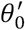 is to differentiate it from their counterparts in (i). If *β* is significantly different from zero, then the mediator has an effect on the outcome after controlling for the independent variable and covariates.

The univariate variable *M* is said to significantly mediate the relationship between *X* and *Y*, if both *α* and *β* are significantly different from zero after accounting for the covariates. Said in a different way, if the product *αβ* is non-zero, then *M* is a mediator for *X* and *Y*, and the product *αβ* quantifies the mediation effect. In the language of a graphical model, this means that both *α* and *β* edges in **Figure** 2 (a) exist, connecting nodes *X* and *Y* via a pathway passing through node *M*.

### Multivariate mediation analysis

Mediation analysis concerning a multivariate mediator can be conducted using structural equation models (SEMs)^6,8^ (see **Figure 2** (b)). Formally, consider *V* mediators (*V* ≥ 2), denoted as *M*(1), *M*(2), *…, M*(*V*), an independent variable *X*, and an outcome variable *Y*. Multivariate mediation analysis considers two conditional models as follows.

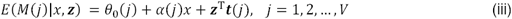

where *M*(*j*) is the *j*^*th*^ mediator; *θ*_0_(*j*), *α*(*j*), and ***t***(*j*) are coefficients for the intercept, the observed independent variable *x*, and the observed covariates ***z*** that are associated with the *j*^*th*^ mediator.

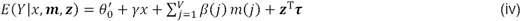

where ***M*** = (*M*(1), *M*(2), *…, M*(*V*)) is a vector representing *V* mediators; 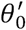, *γ*, and **τ** are coefficients for the intercept, the observed independent variable *x*, and the observed covariates ***z***; *β*(*j*) is the coefficient associated with the *j*^*th*^ mediator.

The *j*^*th*^ mediator *M*(*j*), for *j* = 1, 2, …, *V*, is said to significantly mediate the relationship between *X* and *Y*, if both *α*(*j*) and *β*(*j*) are significantly different from zero after accounting for the covariates. The product, or *α*(*j*)*β*(*j*), quantifies the mediation effect for the *j*^*th*^ mediator (see **Figure 3**).

**Figure 3.**
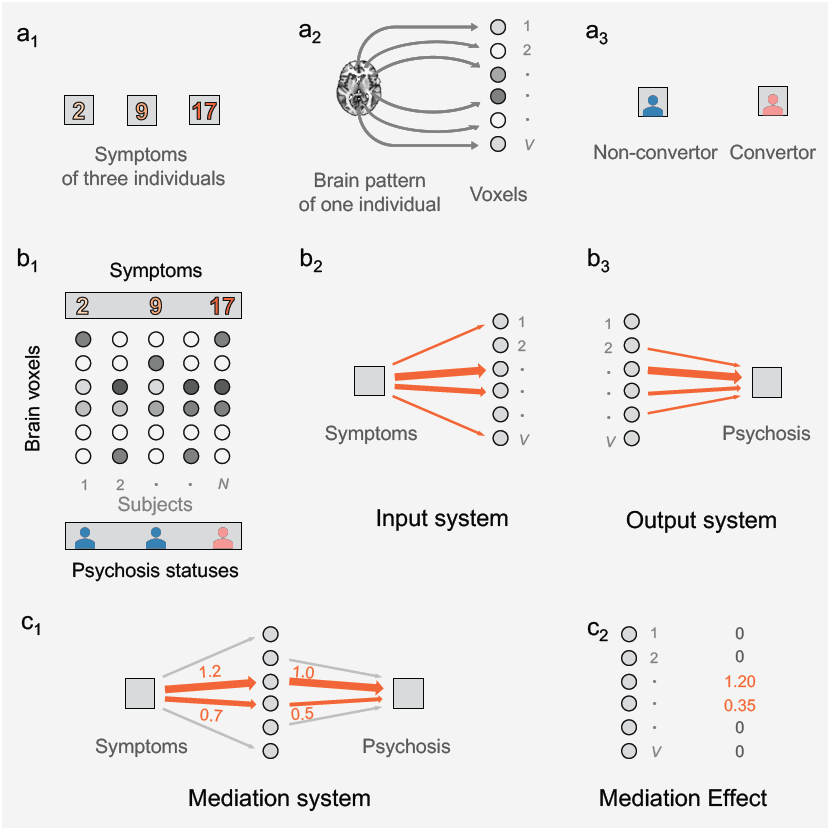
A hypothetical experiment and how multivariate mediation analysis can be used to study brain mediation in health and disease. (a1) Three individuals’ behavioral symptom scores are measured at baseline. (a2) The fMRI BOLD time series for one individual is collapsed into the pattern of activity across voxels in the whole brain averaged over time. Each circle corresponds to one voxel. (a3) Individuals’ two-year clinical outcome for psychosis. We use 0 to refer to a non-convertor and 1 for a convertor. (b_1_) Individuals’ brain patterns are arranged corresponding to their behavioral symptom scores and psychosis statuses. Each column contains data from a particular subject. (b_2_) The input system of the brain mediation framework studies the association between the behavioral symptom score (the box) and brain patterns across mediators (the circles). The pink arrows indicate pathways from the behavioral symptom score to the mediators. The width of the arrows indicates effect size. (b_3_) The output system of the brain mediation framework studies the association between the brain patterns across mediators (the circles) and disease status (the box). (c1) The mediation analysis framework combines the input and output systems, and studies how the effect of behavioral symptom score (the left box) on the psychosis status (the right box) is intermediated by patterns of the neural mediators (the circles). A voxel significantly mediates the relationship if its pattern is associated with both the behavioral symptom score and the psychosis status. (c_2_) The mediation effect of a particular voxel is calculated by multiplying the effect sizes from the input and the output pathway corresponding to the voxel.

### High-dimensional brain-wide functional mediation

High-dimensional mediation analysis aims at identifying mediators from a high-dimensional multivariate variable. For example, a neurobiologist is interested in searching through the entire brain to look for neural mediators using fMRI BOLD signals recorded from hundreds of thousands of brain regions. The high-dimensional functional mediation framework introduced in the paper consists of a dual system: the input system investigates how an independent variable (*e.g*., behavioral symptoms) affects brain patterns, after controlling for covariates; the output system examines how brain patterns give rise to the outcome variable (*e.g*., psychosis disease status), after controlling for the independent variable and covariates (see **Figure** 3). In the following, we introduce the key concepts of the framework and leave derivations and discussions to the **Methods** and **Supplementary Information** sections.

Consider ***N*** subjects and *V* brain areas, where *V* is high-dimensional (in our study *V* = 130,992). Let *x*_*i*_ and *y*_*i*_ be the independent and outcome variables for subject *i*, respectively. Let ***z***_*i*_ = (*z*_1*i*_, *z*_2*i*_, *z*_3*i*_) denote the covariates of subject *i*, for example, the site (from which data are collected), age, and gender, respectively. Finally, let *m*_*ij*_ be the neural activity from the *j*^*th*^ brain area of subject *i*, The high-dimensional functional brain-wide mediation framework consists of a dual functional system: an input system and an output system.

The input system (see **Figure** 3 b_2_) consists of the following conditional model:

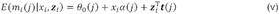

where *θ*_0_(*j*), *α*(*j*), and ***t***(*j*) are coefficients for the intercept, the independent variable, and covariates that are associated with the *j*^*th*^ mediator.

The output system (see **Figure** 3 b_3_) is represented by the following functional model:

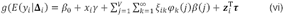

where *β*_0_, *γ*, and **τ** are coefficients for the intercept, the independent variable, and covariates. ξ_*ik*_ and φ_*k*_(*j*) are the Karhunen-Loève expansion^18,19^ of *m*_*ij*_, the *j*^*th*^ mediator of subject *i*; in other words 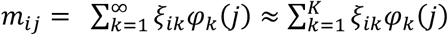 (without loss of generality, assume *m*_*ij*_ is zero centered), where *ξ*_*ik*_*∼****N***(0, *λ*_*k*_), *λ*_1_ *≥ λ*_2_ *≥* … *≥ λ*_*∞*_, for *i* ∈ {1, 2, …, ***N***}, *j* ∈ {1, 2, …, *V*}, and *K* is a finite integer such that 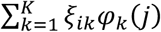 captures the importance of modes of variations of *m*_*ij*_ (see **Methods** for more detailed explanation). ***φ*** = {*φ*_1_, *φ*_2_, …, *φ*_*∞*_} denotes the basis functions. *β*(*j*) is the coefficient associated with the *j*^*th*^ mediator. The link function *g*(·) takes various forms based on the outcome distribution. For example, when *y*_*i*_ is Gaussian, binary, or Poisson, the link function is 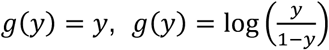, or *g*(*y*) = *log* (*y*), respectively.

The first *K* basis functions or {*φ*_1_, *φ*_2_, …, *φ*_*K*_} in Equation (vi) are functional representations of *K* dominant population-specific (*i.e*., shared by all ***N*** subjects) patterns of brain data, ranked decreasingly (according to {*λ*_1_, *λ*_2_, …, *λ*_*K*_}) based on the amount of information each basis function explains about the neural mediator ***M***. Although by Mercer’s theorem^Chapter 4 of 20^, the basis functions are orthogonal, and researchers indeed are oftentimes interested in uncovering orthogonal signals of brain patterns in relation to mediation, it remains possible that the brain patterns consist of non-orthogonal signals. Despite the central focus herein being on mediation studies, it is important to understand how the underlying orthogonality of brain data’s basis functions may affect the identification of neural mediators and the estimation of the mediation effect. To that end, we performed simulation studies under different noise levels and sample sizes, and the results showed that our framework was successful to uncover neural mediators regardless of the orthogonality of basis functions. Naturally, the mediation analysis performance was better when the underlying basis functions were orthogonal, and the estimation results improved as the noise decreased or the sample size increased (see **Supporting Information**).

The key steps of the framework can be summarized as follows. First, it estimates the effect of the independent variable (*e.g*., the difference between the prodromal *+* and prodromal *−* group) on each brain area, after controlling for all covariates; this yields an ***α***brain atlas (see **Figure 3** (b_2_)). Next, it extracts subject-specific principal component (PC) scores or eigen-values ***ξ***_*i*_ = {*ξ*_*i*1_, *ξ*_*i*2_, …, *ξ*_*iK*_} from each individual *i*’s brain patterns (***m***_*i*_), and estimates the effect of the transformed lower-dimensional mediators (*i.e*. ***ξ***_*i*_) on the disease outcome *y*_*i*_ controlling for the independent variable and the covariates. Subsequently, the low-dimensional mediator-on-outcome effect is translated to the high-dimensional brain space using the estimated brain-wide basis functions of ***φ***; this produces a ***β*** brain atlas (see **Methods** and **Supporting Information**). Finally, it obtains the brain-wide mediation effect using the ***α***and ***β*** brain atlases and bootstrap tests (see **Figure 3** (c_1_) - (c_2_) and **Methods**).

We evaluated the efficacy and utility of the framework using both simulated (see **Supporting Information**) and empirical data from the second phase of the North American Prodrome Longitudinal Study (NAPLS-2^21^) (see **Results**). During simulation studies, we considered 12 scenarios of data generating mechanisms, consisting of different levels of sample sizes, noise, and both orthogonal and nonorthogonal basis functions. The simulation results showed that the framework performed well across different scenarios; the performance improved when the sample size increased or when the noise level decreased. For the empirical data, we aimed at identifying brain regions that mediated individuals’ psychotic symptoms and clinical outcomes in a sample of 263 subjects at clinical high risk (CHR) for psychosis, among whom 25 subjects developed a full-blown psychotic disorder during a 2-year clinical follow-up (CHR convertors). Our results unveiled two sets of neural mediators: the *P* mediators and the *N* mediators. The *P* mediators contained brain areas that positively mediated higher prodromal symptoms and psychotic conversion; the *N* mediators consisted of brain regions that negatively mediated higher prodromal symptoms and psychotic conversion (suggesting a potential protective effect).

## RESULTS

We first conducted simulation studies to ensure that the proposed framework was able to identify brain areas that intermediate the treatment and the outcome under different settings (**Supporting Information**).

After verifying the performance of the proposed framework using simulated data, we applied the framework to an empirical study to identify and isolate functional brain regions that mediate prodromal symptoms at baseline and two-year clinical outcome in subjects at clinical high risk (CHR) for psychosis. The sample included 263 subjects recruited from eight study sites across the United States and Canada who met criteria for a prodromal risk syndrome^22^ at the point of recruitment and were clinically followed up for two years as part of the NAPLS-2 project^21^. During the follow-up period, 25 subjects developed a full-blown psychotic disorder (CHR convertors); 238 did not (CHR non-convertors). All participants received an eyes-open resting-state functional magnetic resonance imaging (fMRI) scan at the point of recruitment.

After data preprocessing, the time series for each voxel within a binary whole-brain mask (130,992 voxels in total) were extracted. These time series were further corrected for physiological and head motion noise and were temporally filtered (bandpass 0.008-0.1 Hz). The prodromal symptoms were quantified using the Scale of Prodromal Symptoms^23^, and the clinical outcome was labeled as convertor or non-convertor.

Since disorganization symptoms have been shown to be a potential clinical predictor for psychosis^1–3^, we first investigated whether such measure was significantly different between convertors and non-convertors at baseline. Using the Welch two sample *t*-test and Pearson’s (product moment) correlation coefficient test, the data confirmed that the convertors and non-convertors in the study indeed had significantly different behavioral symptom scores (*t* = 3.49, *P* < 0.005; Pearson correlation *r* = *−*0.22, *P* < 0.001) (see **Figure** 2 b). We then continued to investigate which brain regions would functionally mediate this association. First, we tested if behavioral symptoms had an effect on any of the 130,992 voxels, controlling for covariates (see **Figure 3** (b_2_)). This analysis yielded the ***α***brain atlas (see **Figure** 4 (a)); each of its 130,992 elements indicated the effect of behavioral symptoms on a brain voxel, after controlling for covariates. Second, we tested if activity from a brain voxel would increase (or decrease) the likelihood of developing psychosis, while controlling for behavioral symptoms and covariates (see **Figure 3** (b_3_)). This was conducted in a generalized principal component estimation model (see **Methods**). This analysis yielded the ***β*** brain atlas (see **Figure** 4 (b)); each of its 130,992 elements represented the effect from a brain voxel to the likelihood of developing psychosis, controlling behavioral symptoms and covariates. Third, we obtained the brain-wide functional mediators using the ***α*** and ***β*** brain atlases and classified them into two categories: the *P* neural mediators and the *N* neural mediators (see **Figure 3** (c_1_) - (c_2_)). Finally, we conducted bootstrap experiments to test whether the mediation effect of each voxel was statistically significant (see **Methods**). All steps had included site (from which data were collected), gender, and age as covariates to remove their confounding effects. Results were reported after Bonferroni correction across all voxels in the brain (see **Figure 4**).

**Figure 4.**
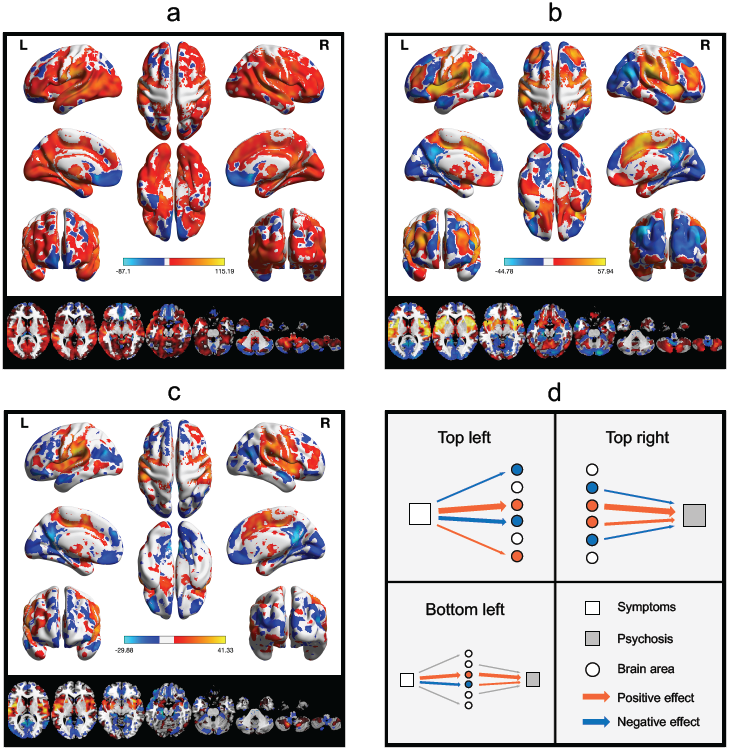
Brain areas that mediate behavioral symptoms and the development of psychosis. A high-dimensional mediation analysis on 130,992 brain voxels of 263 subjects demonstrates that the pathway between behavioral symptoms and psychosis is positively mediated by the right lateral prefrontal cortices, bilateral insular and opercular areas, bilateral sensorimotor areas, striatum, and cerebellar lobules 4, 5, and 6, and negatively mediated by the bilateral medial frontal and orbitofrontal cortices, left lateral prefrontal cortices, posterior cingulate, precuneus, visual cortex, and cerebellar Crus 1 and lobule 9. (a) The ***α*** brain atlas shows surface and subcortex areas associated with behavioral symptoms when controlled for covariates. (b) The ***β*** brain atlas shows surface and subcortex areas associated with psychosis status when controlled for behavioral symptoms and covariates. (c) The *P* and *N* neural mediators. The neural mediators include surface and subcortex areas that are jointly associated with behavioral symptoms and brain disease status, after all covariates are controlled. The areas highlighted in red are *P* mediators, and those highlighted in blue represent *N* mediators. (d) Explanations regarding the pathways and color codes of figures (a) - (c). The color bars indicate effect sizes from bootstrap experiments.

### The *α* and *β* brain atlases

We further inquired into the ***α*** brain atlas (**Figure** 4 (a)) and the ***β*** brain atlas individually (**Figure** 4 (b)) in order to investigate how the input and output systems contribute to overall mediation. Specifically, the ***α*** brain atlas included brain regions that were associated with behavioral symptoms when controlled for covariates; the ***β*** brain atlas consisted of brain areas that were associated with psychosis status when controlled for behavioral symptoms and covariates. In the ***α*** brain atlas, activities of the majority of brain regions were positively associated with behavioral symptoms, while activities of the orbitofrontal cortex and cerebellar crus 1, crus 2, and vermis were negatively correlated with behavioral symptoms. In the ***β*** brain atlas, positive associations were present in the lateral prefrontal cortex, sensorimotor cortex, insular and opercular areas, anterior cingulate cortex, striatum, and cerebellar lobule 9, crus 1, and crus 2. In contrast, negative associations were shown in the medial prefrontal cortex, posterior cingulate cortex, visual cortex, and cerebellar lobules 6, 7b, and 8.

### The *P* and *N* mediators

Discovering brain areas that are positively and negatively mediating behavioral symptoms and disease development is a central problem in neuropathology. To promote discussion, here we defined the *P* mediators as brain areas whose activities were positively mediating higher behavioral symptoms and increased chance of conversion to psychosis (see regions with positive weights in see **Figure 4** (c)). In other words, they are potentially facilitative of conversion to psychosis among subjects with higher behavioral symptoms. Our results revealed that the *P* mediators were mainly present bilaterally at the right lateral prefrontal cortices, bilateral insular and opercular areas, bilateral sensorimotor cortices, striatum, and cerebellar lobules 4, 5, and 6. In contrast, we defined the *N* mediators as brain areas whose activities were negatively mediating higher behavioral symptoms and increased chance of conversion to psychosis (see regions with negative weights in see **Figure 4** (c)). In other words, they are potentially protective of psychosis conversion among subjects with higher behavioral symptoms. The *N* mediators were located chiefly at the bilateral medial frontal and orbitofrontal cortices, left lateral frontal cortex, posterior cingulate, precuneus, visual cortex, and cerebellar crus 1 and lobule 9.

## DISCUSSION

In this study, we designed a brain-wide functional mediation analysis framework, and, through its lenses, identified and isolated *P* and *N* neural mediators that mediate psychosis prodromal behavioral symptoms and disease status among individuals at CHR and quantified each mediator’ effects on developing psychosis. The *P* mediators consisted of neural signatures associated with positive mediation, which were primarily distributed in the brain’s sensorimotor system, insular and opercular areas, and striatum; the *N* mediators consisted of neural signatures associated with negative mediation, which were chiefly located in the brain’s default-mode system and visual system. The isolation of the *P*-*N* mediators highlights the neurobiological pathways from early psychotic signs to full-blown psychosis and their identification demonstrates the utility of the proposed methodological framework in clinical neuroscience studies.

The proposed framework showed promise to study how brain patterns may intermediate between an independent variable (such as prodromal behavioral symptoms) and an outcome variable (such as developing psychosis). Using a generalized principal component estimation model and functional data analysis, the framework extends multivariate mediation analysis to the terrain of high-dimensional functional mediation analysis with non-Gaussian outcomes. To inquire into the functional organization of the mediator, the model extracts subject-specific principal component (PC) scores and functional representations (population-specific brain-wide basis functions) of brain patterns. The estimated effect from PC scores to psychosis status is then translated to the whole brain space via the brain-wide basis functions. A *logit* link function was employed to couple patterns of the brain and the independent variable with the disease outcome. Since the model allows for a variety of link functions, it can assist other mediation problems with outcome distributions from the exponential family.

There are a few additional properties of the proposed framework that may be useful in other studies. First, the framework integrates a generalized principal component estimation model into a dual regulatory system connecting an input system and an outcome system. This technique may shed some light on how to build biological architecture consisting of sub-systems. Second, when the outcomes are binary, one can use the framework to evaluate controlled direct effects, and natural direct and indirect effects on the odds-ratio scale (**Supporting Information**). Third, high-dimensional brain patterns may contain multilevel information. This framework allows us to extract both group-level (*i.e*., the group-level basis functions) and subject-specific features (*i.e*., the subject-specific PC scores) of the brain data. The subject-specific features may be used as compact neural signatures to assess individual differences in the future (**Supporting Information**).

We applied our proposed framework in a psychosis neuroimaging study, where we investigated how functional brain activities may mediate prodromal behavioral symptoms and clinical outcome. The findings observed in this study confirmed previous studies and provided additional, useful insights for future psychosis research. First, our findings suggest that the *P* mediators are largely distributed in the brain’s primary functional systems (*e.g*., sensorimotor cortex, cerebellar lobules 4, 5, 6, insular-opercular area including the superior temporal cortex and Heschl’s gyrus). These regions are involved in basic sensory functions in humans such as perception, hearing, and motion. A large number of studies have demonstrated that increased activities in the insular-opercular area are strongly associated with auditory hallucinations in patients with schizophrenia^24–26^, and increased sensorimotor connectivity is robustly present across the entire course of schizophrenia from prodromal phase to chronic patients^27–29^. The exact mechanisms underlying the hyperactivity state of these sensory systems are, however, unclear; they have been putatively considered as either a result of sensory gating deficits disrupted by excessive mesolimbic dopamine input^30^ or as reflective of aberrant top-down cognitive control associated with strong perceptual priors^26^. In line with previous findings, the current study further shows that such increased activities may be potential mediators between prodromal symptoms and psychosis conversion, pointing to a critical role of sensory systems in the development of psychotic disorders. Second, the *N* mediators are primarily distributed in the brain’s default-mode network (DMN) and limbic system, including medial frontal cortex, orbitofrontal cortex, posterior cingulate cortex, precuneus, and cerebellar crus 1. The DMN is one of the most frequently reported systems whose function is strongly associated with psychosis. The most prominent finding regarding DMN in patients is the failure to deactivate this network during cognitive tasks^31–33^, which may relate to exaggerated internally-focused thoughts and lack of sufficient suppression of these thoughts during cognition^34^. Here, the finding of negative mediation effect in DMN activities during resting state is parallel to such interpretation, suggesting lower activity (indicating insufficient activation) during rest may potentially mediate prodromal symptoms and psychosis status. In contrast, higher DMN activity during rest may serve as a protector against conversion.

A few reasons have made the blood-oxygen-level-dependent functional magnetic resonance imaging (BOLD fMRI) data suitable for studying brain-wide mediation. First, although studies have used resting state electroencephalography (EEG) data and discovered brain areas, such as the frontal regions, that are associated with psychosis^35^, imaging modalities with greater spatial resolution, such as fMRI, may both confirm and extend neural signatures beyond those identified using EEG. Second, reduced auditory P300 event-related potential (ERP) amplitude (from a functional neurophysiological test) is a primary candidate electrophysiologic biomarker of psychosis^36^; it nevertheless may not capture as much variability that occurs in spontaneous brain activity as the fMRI data to work well as a biomarker for conversion. Third, slow wave power has been shown to correlate with reduced blood flow and glucose utilization in schizophrenia patients, and is therefore thought to reflect reduced functioning in the frontal area^37,38^. This supports the utility of fMRI BOLD data in mediation studies. Finally, structural MRI studies are beginning to discover associations between structural brain information and conversion to psychosis^9^; here we have shown that functional MRI data could add new insights into studies of neural markers associated with psychosis.

Although we demonstrated high-dimensional functional neural mediation analysis in the domain of brain studies, the framework may also be useful to study other high-dimensional mediation problems, such as how genome-wide genotypes mediate the effect of environmental factors on phenotypes. With the recent convergence in neuroimaging, genomics, health informatics, wearable and digital sensors, the model has the potential to identify a broad range of novel, informative, and previously unavailable biomarkers that play an important role in mediating the relationship between healthy and diseased agents and their environment. For example, the model may be used to understand how gene expression, brain physiology, and circadian patterns jointly mediate environment and biological phenotypes, uncover brain regions that mediate sensory input and behavior outcome collected by wearable devices, and study how computers can act as a mediating artificial intelligence (AI), transferring human input into computer-generated intelligent responses.

There are several limitations with the proposed method. First, in clinical practice, one assumes that changing brain patterns can first cause prodromal signs and symptoms, followed in some cases by later conversion to psychosis. In this paper, we aimed at studying the influence of the underlying brain patterns, which were not directly observable, on the link between two directly and clinically observable sets of variables: prodromal signs and symptoms on the one hand, and conversion state on the other hand. The framework we designed to map the pathways contained directed arrows that clarified the statistical model was mediation analysis and did not suggest a definitive causal flow from prodromal signs via brain patterns towards conversion status (see **Figures** 1 and 2). When confusion about the causal direction existed, one can interpret the identified neural mediators as brain areas that are jointly associated with behavioral symptoms and psychosis conversion. In other words, the mediators exclude brain areas that are associated with conversion, but that are not associated with prodromal symptoms, and *vice versa*. Second, we were mainly interested in identifying brain regions that were simultaneously mediating behavioral symptoms and psychosis status. This naturally left out the cases where some mediators were interposed before or after other mediators. Future analysis may incorporate dynamic mediating systems and information feedback components. Third, throughout, we assumed that behavioral symptoms did not interact with the mediators. Future work that includes interaction between the independent variable and the mediator may be useful to expand current analysis (see **Supporting Information** for an example and see^39^ for a special case). Fourth, although dimension reduction could reduce biases caused by spurious correlation, our framework cannot remove the coincidental association between some (voxel) features and the residual term (*i.e*., incidental endogeneity). This is an active research area in high-dimensional data analysis^see 40^. Fifth, the proposed model focused on neural markers by averaging the brain time courses over time. This omitted the territory of mediation analysis where the mediation effect changes over time. Future work needs to extend the framework to longitudinal settings: such extensions are particularly useful for studying causal mediation effect related to brain development during childhood and adolescence, brain aging between health and disease, and brain degeneration along the trajectory of a neurodegenerative disease development. We are currently investigating how to extend the techniques used in our framework to improve our understanding about large-scale longitudinal mediation, such as combing functional data analysis (FDA) and dual mediation system with autoregressive models^41,42^, latent growth curve (LGM)^43^, parallel process models^44^, latent difference score (LDS) models^45^, and autoregressive LGM models^46^.

To summarize, in the present study, we propose a framework that leverages statistical- and machine-learning algorithms for neuroscientific insights into understanding the properties of large-scale functional intermediating neural markers that mediate the relationship between an independent variable and a binary disease outcome. The conceptual and analytical architecture of the framework in dealing with high-dimensional data and its flexibility in handling non-normally distributed outcomes have the potential to be widely adapted to diverse scenarios of causal paradigms to uncover intermediating pathways in complex regulative systems.

## METHODS

### The NAPLS-2 Data

The fMRI data were drawn from the second phase of the North American Prodrome Logitudinal Study (NAPLS-2) consortium^21^, which included 263 subjects recruited from eight study sites across the United States and Canada. All subjects met the criteria for the prodromal syndromes at the point of recruitment according to the Structured Interview for Prodromal Syndromes (SIPS, ^23^) and were clinically followed-up for two years. During follow-up, 25 subjects developed one type of the Axis-I psychotic disorders (CHR convertors, age 18.52 ± 4.08 years, 17 males) and 238 did not (CHR non-convertors, 19.07 ± 4.16 years, 136 males). All subjects received an eyes-open resting-state functional magnetic resonance imaging (fMRI) scan at the point of recruitment.

### Data Acquisition

During the 5-min eyes-open resting-state scan (154 whole-brain volumes), participants were asked to lay still in the scanner, relax, gaze at a fixation cross, and not engage in any particular mental activity. After the scan, investigators confirmed with the participants that they had not fallen asleep in the scanner. Data were acquired from 3T MR scanners located at eight study sites with identical fMRI protocols. Siemens scanners were used at Emory, Harvard, University of California Los Angeles (UCLA), University of North Carolina at Chapel Hill (UNC) and Yale, and GE scanners were used at Calgary, University of California San Diego (UCSD) and Zucker Hillside Hospital (ZHH). Functional images were collected using gradient-recalled-echo echo-planar imaging (GRE-EPI) sequences: TR/TE 2000/30 ms, 77 degree flip angle, 30 4-mm slices, 1-mm gap, 220-mm FOV. In addition, we also acquired high-resolution T1-weighted images for each participant with the following sequence: 1) Siemens scanners: magnetization-prepared rapid acquisition gradient-echo (MPRAGE) sequence with 256 mm x 240 mm x 176 mm FOV, TR/TE 2300/2.91 ms, 9 degree flip angle; 2) GE scanners: spoiled gradient recalledecho (SPGR) sequence with 260 mm FOV, TR/TE 7.0/minimum full ms, 8 degree flip angle.

### Preprocessing of rs-fMRI Data

Data were preprocessed using the standard pipeline implemented in the Statistical Parametric Mapping (SPM12, http://www.fil.ion.ucl.ac.uk/spm/) software following previously published work^47–50^, including slice-timing correction, realignment, individual structural-functional image coregistration, normalization to the Montreal Neurological Institute (MNI) template and spatial smoothing with 8-mm full width at half maximum (FWHM). Preprocessed fMRI time series were extracted from a total of 130,992 voxels covering the entire brain for each subject. The extracted time series were corrected for white matter and cerebrospinal signals, 24 head motion parameters (6 translation and rotation parameters, their first derivatives, and the square of these 12 parameters), and frame-wise displacement, and were then temporally filtered (band-pass 0.008-0.1 Hz). Secondary data analysis was conducted using the R software.

### The Model

We designed a dual system for high-dimensional functional mediation analysis. The input system tests if the independent variable (*e.g*. behavioral symptoms) has an effect on each mediator, after controlling for all covariates (see **Figure 3** (b_2_)). The output system uses a generalized principal component estimation model to test if each mediator has an effect on the outcome, after controlling for the independent variable and covariates (see **Figure 3** (b_3_)).

Formally, consider ***N*** subjects and *V* brain areas, where *V* = 130,992. Let *x*_*i*_ ∈ ℝ be the independent variable for subject *i*, let covariates ***z***_*i*_ ∈ ℝ^3^ consists of the site (from which data are collected), age, and gender, respectively, of subject *i*, ***m***_*i*_ ∈ ℝ^*V*^ be the brain patterns of subject *i* spanning *V* brain areas, and *y*_*i*_ be the outcome for subject *i* (in this study *y*_*i*_ ∈ {0,1}).

We first review the basics of functional principal component analysis. Let *m*(*j*), *j* ∈ [0,1], be a squared integrable random function with mean *μ*(*j*) and covariance function *K*(*s, t*). In other words, *μ*(*j*) = *E*(*m*(*j*)) and *K*(*s, t*) = *cov*(*m*(*s*), *m*(*t*)). By Mercer’s theorem^Chapter 4 of 20^, one can obtain the spectral decomposition of *K*(*s, t*) as:

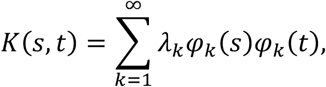

where *λ*_1_ *≥ λ*_2_ *≥* … *≥* λ_*∞*_ are decreasingly ordered nonnegative eigenvalues and φ_*k*_’s are their corresponding orthogonal eigenfunctions with unit ℒ^2^ norms.

Karhunen-Loève expansion^18, 19^ of the random function *m*(*j*) yields:

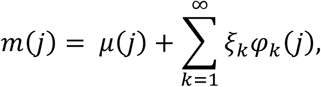

where 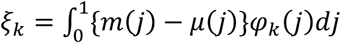 are uncorrelated random variables with zero mean and variance λ_*k*_. For a given functional sample, the mean function *μ*(*j*) and covariance function *K*(*s, t*) can be consistently estimated using the method of moments. The eigen -values and -functions are estimated from the empirical covariance function, and the principal component scores (*ξ*_*k*_’s) can be estimated by numeric integration.

Next, we enquire further into Equations (v) and (vi) from the **Introduction** section.

The input system consists of the following model:

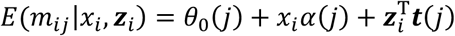

*θ*_0_(*j*), *α*(*j*), and ***t***(*j*) are coefficients for the intercept, the independent variable, and covariates that are associated with the *j*^*th*^ mediator. Specifically, *α*(*j*) captures the effect of the independent variable on the *j*^*th*^ mediator, *θ*_0_(*j*) indicates an intercept, and ***t***(*j*) denotes the coefficients for the covariates with respect to the *j*^*th*^ mediator. Without loss of generality, consider 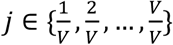.

The output system consists of the following functional model:

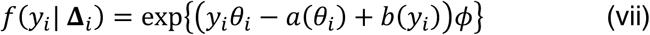

where *θ*_*i*_ = ***h***(*η*_*i*_), *η*_*i*_ is called a linear predictor with 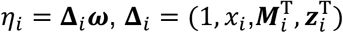 denotes the data, 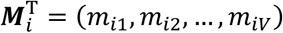, and 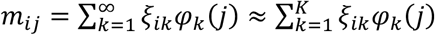, where ξ_*ik*_*∼****N***(0, λ_*k*_), *λ*_1_ *≥ λ*_2_ *≥* … *≥ λ*_*∞*_, for *i* ∈ {1, 2, …, ***N***} and 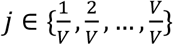, and ***φ*** = {*φ*_1_, *φ*_2_, *…, φ*_*K*_} is a set of basis functions. ***ω*** = (*β*_0_, *γ*, ***β*, τ**)^T^ denotes the corresponding parameters for **Δ**_*i*_. ***h***(·), *a*(·) and *b*(·) are proper functions. ***β*** is a *V* × 1 vector, whose *j*^*th*^ entry *β*(*j*) estimates the effect of the *j*^*th*^ mediator on the outcome, controlling the independent variable and covariates. *ϕ* is a nuisance parameter.

By taking the expected value of *y*_*i*_ conditioning on **Δ**_*i*_, Equation (vii) yields Equation (vi) in the **Introduction** section. Particularly, when *y*_*i*_ is binary, (vi) has the following form:

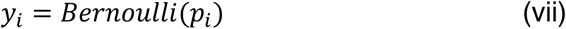

where 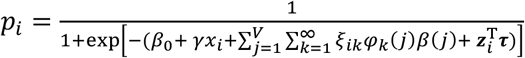

Thanks to spectral decomposition of ***M***, parameter estimation of ***β*** can be performed on ***ξ*** = (***ξ***_1_, ***ξ***_2_, *…*, ***ξ***_*N*_)^T^, where ***ξ***_*i*_ = {*ξ*_*i*1_, *ξ*_*i*2_, …, *ξ*_*iK*_}. Toseethis,rewrite 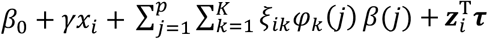 in (vii) as 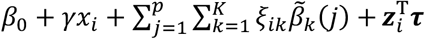, where 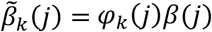 The estimation problem now translates to estimating the low-dimensional mediator-on-outcome effect, or 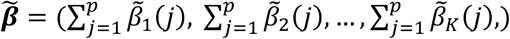 Subsequently, the estimation of ***β*** can be retrieved by projected the estimated 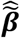 back to the brain space using the estimated basis functions 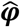; in other words, 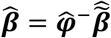.

Although in real world brain data, the (unknown) basis functions ***φ*** can be either orthogonal or non-orthogonal, simulation studies showed that, regardless of the orthogonality of basis functions, our framework was successful to uncover the ***α*** and ***β*** brain atlases under different noise levels and sample sizes (**Supporting Information**).

Since ***N****IE*(*j*) = *α*(*j*)*β*(*j*) has a one-to-one relationship to the *j*^*th*^ voxel’s mediation effect (**Supporting Information**), for simplicity we estimate ***N****IE*(*j*), for *j* = 1, 2, …, *V*. In words, ***N****IE*(*j*) is the *j*^*th*^ voxel’s natural indirect mediation effect on the *log* odds-ratio scale per unit increase of the independent variable. When *V* is small, Sobel’s Test^51^ evaluates the statistical significance of ***N****IE*(*j*); when *V* is high-dimensional, the statistical significance of ***N****IE*(*j*) can be evaluated using a bootstrap approach^52^ (**Supporting Information**); the results are further adjusted for multiple comparisons.

### Brain-wide functional mediation analysis

We conducted brain-wide functional mediation analysis on the rs-fMRI data from the NAPLS-2 sample using the proposed framework. We first assessed if there was any effect from the independent variable on each voxel, controlling for all covariates, using Equation (v) and obtained the ***α*** brain atlas. We then estimated the effect each mediator had on the outcome, controlling for the independent variable and all covariates, using Equation (vi) and obtained the ***β*** brain atlas. Finally, we identified *P* neural mediators and *N* neural mediators and estimated their mediation effect via bootstrap experiments. We presented the empirical results in **Figure 4**.

## Author contributions

O.Y.C. designed the model and performed the mediation analysis. H.C. provided neurobiological intepretations. G.N. and J.M.R. provided neurobiological support. H.P. and J.P. provided machine-learning support. J.G., T.Q., and J.D. provided statistical support. T.D.C. and M.D.V. provided funding, support, and guidance. O.Y.C. and H.C. wrote the manuscript, with comments from all other authors.

## Acknowledgement

The research was supported by the Wellcome Trust SCNI (098461/Z/12/Z) to Dr. de Vos, the NARSAD Young Investigator Grant (No. 27068) to Dr. Cao, and the NIH grant U01 MH081902 to Dr. Cannon. T.C. would like to acknowledge gifts from the Staglin Music Festival for Mental Health and International Mental Health Research Organization. The authors wish to thank the principal investigators of the North American Prodrome Longitudinal Study for allowing use of the project’s imaging data in this demonstration analysis: Jean Addington, Carrie Bearden, Kristin Cadenhead, Barbara Cornblatt, Diana Perkins, Larry Seidman, Elaine Walker, and Scott Woods.

## Competing interests

The authors declare no competing interests.

## Notes

### Competing Interest Statement

The authors have declared no competing interest.

## References

1. Cannon, T. D. et al. Prediction of Psychosis in Youth at High Clinical Risk. Arch. Gen. Psychiatry 65, 28–37 (2008).

2. Demjaha, A., Valmaggia, L., Stahl, D., Byrne, M. & McGuire, P. Disorganization/cognitive and negative symptom dimensions in the at-risk mental state predict subsequent transition to psychosis. Schizophr. Bull. 38, 351–359 (2012).

3. Carrión, R. E. et al. Prediction of functional outcome in individuals at clinical high risk for psychosis. JAMA Psychiatry 70, 1133–1142 (2013).

4. Chén, O. Y. et al. High-dimensional multivariate mediation with application to neuroimaging data. Biostatistics 19, 121–136 (2018).

5. Geuter, S. et al. Multiple brain networks mediating stimulus-pain relationships in humans. Cerebral Cortex bhaa048 (2018).

6. VanderWeele, T. J. & Vansteelandt, S. Mediation Analysis with Multiple Mediators. Epidemiol. Method. 2, 95–115 (2014).

7. Huang, Y. T. & Pan, W. C. Hypothesis test of mediation effect in causal mediation model with high-dimensional continuous mediators. Biometrics 72, 402–413 (2016).

8. Lindquist, M. A. Functional causal mediation analysis with an application to brain connectivity. J. Am. Stat. Assoc. 107, 1297–1309 (2012).

9. Chung, Y. et al. Adding a neuroanatomical biomarker to an individualized risk calculator for psychosis: A proof-of-concept study. Schizophr. Res. 208, 41–43. (2019).

10. Baron, R. M. & Kenny, D. A. The moderator-mediator variable distinction in social psychological research: conceptual, strategic, and statistical considerations. J. Pers. Soc. Psychol. 51, 1173–1182 (1986).

11. Robins, J. M. & Greenland, S. Identifiability and exchangeability for direct and indirect effects. Epidemiology 3, 143–155 (1992).

12. Whittle, R., Mansell, G., Jellema, P. & van der Windt, D. Applying causal mediation methods to clinical trial data: What can we learn about why our interventions (don’t) work? Eur. J. Pain 21, 614–622 (2017).

13. Fishbein, M. & Ajzen., I. Belief, Attitude, Intention, and Behavior: An Introduction to Theory and Researched. (Addison-Wesley, 1975).

14. Hyman, H. H. Survey Design and Analysis: Principles, Cases and Procedures. (Free Press, 1955).

15. Alwin, D. F. & Hauser, R. M. The Decomposition of Effects in Path Analysis. Am. Sociol. Rev. 40, 37 (1975).

16. Judd, C. M. & Kenny, D. A. Process analysis: Estimating Mediation in Treatment Evaluations. Eval. Rev. 5, 602–619 (1981).

17. Sobel, M. E. Asymptotic confidence intervals for indirect effects in structural equation models. Sociol. Methodol. 13, 290–312 (1982).

18. Karhunen, K. Über lineare Methoden in der Wahrscheinlichkeitsrechnung (On linear methods in probability and statistics). Ann. Acad. Sci. Fenn. Ser. A. I. Math.-Phys. 31, 1–79 (1947).

19. Loève, M. Fonctions aléatoires de second ordre (Random second order functions). In Comptes Rendus De L’Académie Des Sciences 220, (1945).

20. Indritz, J. Methods in Analysis. (Macmillan, 1963).

21. Addington, J. et al. North American Prodrome Longitudinal Study (NAPLS 2): Overview and recruitment. Schizophr. Res. 142, 77–82 (2012).

22. Miller, T. J. et al. Prodromal Assessment with the Structured Interview for Prodromal Syndromes and the Scale of Prodromal Symptoms: Predictive Validity, Interrater Reliability, and Training to Reliability. in Schizophrenia Bulletin 29, 703–715 (2003).

23. McGlashan, T. H., Miller, T. J., Woods, S. W., Hoffman, R. E. & Davidson, L. Instrument for the Assessment of Prodromal Symptoms and States. In Early Intervention in Psychotic Disorders, edited by T. Miller, S.A. Mednick, T.H. McGlashan, J. Libiger, & J.O. Johannessen, pp. 135–149. Springer, Dordrecht (2001).

24. Dierks, T. et al. Activation of Heschl’s gyrus during auditory hallucinations. Neuron 22, 615–621. (1999).

25. Shergill, S. S., Brammer, M. J., Williams, S. C. R., Murray, R. M. & McGuire, P. K. Mapping auditory hallucinations in schizophrenia using functional magnetic resonance imaging. Arch. Gen. Psychiatry 57, 1033–1038 (2000).

26. Powers, A. R., Mathys, C. & Corlett, P. R. Pavlovian conditioning–induced hallucinations result from overweighting of perceptual priors. Science 357, 596–600 (2017).

27. Woodward, N. D., Karbasforoushan, H. & Heckers, S. Thalamocortical dysconnectivity in schizophrenia. Am. J. Psychiatry 169, 1092–1099 (2012).

28. Woodward, N. D. & Heckers, S. Mapping Thalamocortical Functional Connectivity in Chronic and Early Stages of Psychotic Disorders. Biol. Psychiatry 79, 1016–1025 (2016).

29. Anticevic, A. et al. Association of thalamic dysconnectivity and conversion to psychosis in youth and young adults at elevated clinical risk. JAMA Psychiatry 72, 882–891 (2015).

30. Braff, D. L. Sensorimotor Gating and Schizophrenia. Arch. Gen. Psychiatry 47, 181–188 (1990).

31. Pomarol-Clotet, E. et al. Failure to deactivate in the prefrontal cortex in schizophrenia: Dysfunction of the default mode network? Psychol. Med. 38, 1185–1193 (2008).

32. Landin-Romero, R. et al. Failure of deactivation in the default mode network: A trait marker for schizophrenia? Psychol. Med. 45, 1315–1325 (2015).

33. Fryer, S. L. et al. Deficient suppression of default mode regions during working memory in individuals with early psychosis and at clinical high-risk for psychosis. Front. Psychiatry 4, 92 (2013).

34. Whitfield-Gabrieli, S. & Ford, J. M. Default Mode Network Activity and Connectivity in Psychopathology. Annu. Rev. Clin. Psychol. 8, 49–76 (2012).

35. Sollychin, M. et al. Frontal slow wave resting EEG power is higher in individuals at Ultra High Risk for psychosis than in healthy controls but is not associated with negative symptoms or functioning. Schizophr. Res. 208, 293–299 (2019).

36. Hamilton, H. K. et al. Association between P300 Responses to Auditory Oddball Stimuli and Clinical Outcomes in the Psychosis Risk Syndrome. JAMA Psychiatry 76, 1187–1197 (2019).

37. Guich, S. M. et al. Effect of attention on frontal distribution of delta activity and cerebral metabolic rate in schizophrenia. Schizophr. Res. 2, 439–448 (1989).

38. Ingvar, D. H., Sjölund, B. & Ardö, A. Correlation between dominant EEG frequency, cerebral oxygen uptake and blood flow. Electroencephalogr. Clin. Neurophysiol. 41, 268–276 (1976).

39. Muller, D., Judd, C. M. & Yzerbyt, V. Y. When Moderation is Mediation and Mediation is Moderated. J. Pers. Soc. Psychol. 89, 852–863 (2005).

40. Fan, J. & Liao, Y. Endogeneity in high dimensions. Ann. Stat. 42, 872–917 (2014).

41. Gollob, H. F. & Reichardt, C. S. Interpreting and estimating indirect effects assuming time lags really matter. In Best methods for the analysis of change: Recent advances, unanswered questions, future directions, edited by L. M. Collins & J. L. Horn, pp. 243–259. American Psychological Association, Washington, D.C., USA (1991).

42. Cole, D. A. & Maxwell, S. E. Testing Mediational Models with Longitudinal Data: Questions and Tips in the Use of Structural Equation Modeling. Journal of Abnormal Psychology 112, 558–577 (2003).

43. Muthén, B. O. & Curran, P. J. General Longitudinal Modeling of Individual Differences in Experimental Designs: A Latent Variable Framework for Analysis and Power Estimation. Psychol. Methods 2, 371–402 (1997).

44. Cheong, J., Mackinnon, D. P. & Toon, S. Structural Equation Modeling : A Investigation of Mediational Processes Using Parallel Process Latent Growth Curve Modeling. Struct. Equ. Model. A Multidiscip. J. 10, 238–262 (2009).

45. McArdle, J. J. A latent difference score approach to longitudinal dynamic structural analysis. Structural Equation Modeling: Present and Future. In Structural equation modeling: Present and future, edited by R. Cudeck, S. du Toit, & D. Sorbom, pp. 342–380. Scientific Software International, Lincolnwood, IL USA (2001).

46. Bollen, K. A. & Curran, P. J. Autoregressive Latent Trajectory (ALT) Models: A Synthesis of Two Traditions. Sociol. Methods Res. 32, 336–383 (2004).

47. Cao, H. et al. Cerebello-thalamo-cortical hyperconnectivity as a state-independent functional neural signature for psychosis prediction and characterization. Nat. Commun. 9, 3836 (2018).

48. Cao, H. et al. Progressive reconfiguration of resting-state brain networks as psychosis develops: Preliminary results from the North American Prodrome Longitudinal Study (NAPLS) consortium. Schizophr. Res. (2019).

49. Cao, H., Ingvar, M., Hultman, C. & Cannon, T. Evidence for cerebello-thalamo-cortical hyperconnectivity as a heritable trait for schizophrenia. Translational psychiatry 9, 1–8. (2019).

50. Cao, H. et al. Altered Brain Activation During Memory Retrieval Precedes and Predicts Conversion to Psychosis in Individuals at Clinical High Risk. Schizophr. Bull. 45, 924–933 (2019).

51. Sobel, M. E. Asymptotic confidence intervals for indirect effects in structural equation models. Sociol. Methodol. 13, 290–312 (1982).

52. Efron, B. Bootstrap Methods: Another Look at the Jackknife. Ann. Stat. 7, 1–26 (1979).

