## Supplemental Information for "Identifying Neural Signatures Mediating Behavioral Symptoms and Psychosis Onset: High-Dimensional Whole Brain Functional Mediation Analysis"

### 1. Simulation Studies

We simulate data using the following procedures.

Consider $N$ subjects and $V$ brain areas, where $V = 130{,}992$. Let $x_i$ be the independent variable for subject $i$, $z_{i1}$, $z_{i2}$, and $z_{i3}$ be the site (from which data are collected), age, and gender, respectively, of subject $i$, $m_{ij}$ be the neural activity from the $j^{th}$ brain area of subject $i$, and $y_i$ be the outcome for subject $i$.

(1) $x_i = \rho_0 + \rho_1 z_{i1} + \rho_2 z_{i2} + \rho_3 z_{i3} + e_i$

where $z_{i1}$ is a random sample from a set of eight sites (mimicking the distribution of eight sites considered in the study), $z_{i2} \sim \dot{N}(\mu = 19, \sigma = 4.15, l = 12, u = 32)$, where $\dot{N}()$ denotes a truncated normal distribution with mean $\mu$, standard deviation $\sigma$, lower bound $l$ and upper bound $u$, $z_{i3} \sim B(0.42)$. These choices are made to approximate the distribution of covariates used in the real dataset; but can be chosen in other forms without loss of generality. We take parameters $\rho_0 = 6.765$, $\rho_1 = -0.174$, $\rho_2 = -0.072$, and $\rho_3 = -0.456$ to be the estimates from the real data using Equation (1) above. How these parameters are set only affect the simulated values of $x_i$; they will not affect parameter estimation of $\alpha(j)$ and $\beta(j)$ below. Consider noise $e_i \sim N(0, \sigma_e^2)$.

(2) $m_{ij} = \theta_0(j) + \sum_{k=1}^{K} \xi_{ik} \varphi_k(j) + x_i \alpha(j) + z_{i1} t_1(j) + z_{i2} t_2(j) + z_{i3} t_3(j) + \varepsilon_i$

where $\xi_{ik} \sim N(0, \lambda_k)$, $\lambda_k = 0.5^{k-1}$, for $i \in \{1,2,\ldots,N\}$, $j \in \{\frac{1}{V}, \frac{2}{V}, \ldots, \frac{V}{V}\}$, and $k \in \{1,2,\ldots,K\}$. $\{\varphi_1, \varphi_2, \ldots, \varphi_K\}$ is a set of basis functions (see sections 1.1 and 1.2 below). Let $\alpha(j) = \cos(2\pi j)$, $\theta_0(j) = \sin(8\pi j)$, $t_1(j) = \cos(8\pi j)$, $t_2(j) = \sin(12\pi j)$, and $t_3(j) = \cos(12\pi j)$. Consider noise $\varepsilon_i \sim N(0, \sigma_\varepsilon^2)$.

(3) $y_i = Bernoulli(p_i)$, where $p_i = \frac{1}{1+\exp\left[-(\theta_0' + \sum_{j=1}^{V} m_{ij}\beta(j) + x_i\gamma + \tau_1 z_{i1} + \tau_2 z_{i2} + \tau_3 z_{i3} + \epsilon_i)\right]}$.

where $\beta(j) = \sin(2\pi j)$, $\gamma = -0.01$, $\tau_1 = 0.1$, $\tau_2 = 0.1$, and $\tau_3 = 0.1$. Consider noise $\epsilon_i \sim N(0, \sigma_\epsilon^2)$. For each simulation below, $\theta_0'$ is chosen at the value so that the outcomes contain both 0's and 1's (to simulate the scenario where the dataset contain both convertor and non-convertor subjects).

We study different combinations of eigenfunctions ($\{\varphi_1, \varphi_2, \ldots, \varphi_K\}$), idiosyncratic noises ($\sigma_e^2$, $\sigma_\varepsilon^2$, $\sigma_\epsilon^2$), and sample sizes ($N$). For eigenfunctions, we consider $K = 4$ basis functions and study the cases when they are orthogonal and when they are non-orthogonal (to cover the two general scenarios where the brain patterns consist of orthogonal and non-orthogonal neural signals). For noises, we consider a range of magnitudes, at ($\sigma_e^2$, $\sigma_\varepsilon^2$, $\sigma_\epsilon^2$) $= (0.01, 0.01, 0.01)$ (small noise), $(0.1, 0.1, 0.1)$ (moderate noise), and $(5, 5, 5)$ (very large noise). We consider very large noise because we want to see to what degree the estimates would be able to (and unable to) uncover the signals, and whether, even under very large noise level, signals can be recovered when there are more samples. To assess the performance of our method under different sample sizes, we consider $N$ to be 100 and 500. From the estimation performance under the selected combinations of noise levels and sample sizes, one can relatively easily peer into how the model would perform under other magnitudes of noises and samples sizes. Overall, we examine 12 model conditions; for each condition, we conduct 100 bootstrap simulations.

#### A summary of simulation studies

Overall, the framework proposed in this study was able to uncover (simulated) areas of brain involved in mediation. Particularly,

1. The framework successfully uncovered neural mediators across different combinations of eigenfunctions, idiosyncratic noises, and sample sizes.
2. The performance on estimating the input map $\boldsymbol{\alpha}$ was better than it on estimating the output map $\boldsymbol{\beta}$. With larger samples, however, the estimation of $\boldsymbol{\beta}$ improved significantly.
3. Both estimations of $\boldsymbol{\alpha}$ and $\boldsymbol{\beta}$ deteriorated when more noises were added; but for each noise level, the estimation performance improved with more samples. At perhaps the extreme end when the noise was very large, the algorithm was still able to uncover some signals using large samples.
4. The estimation using signals simulated from orthogonal basis functions outperformed those from non-orthogonal basis functions. Both cases saw significant improvement when less noise or larger a sample size was considered.

### 1.1 Orthogonal basis functions

Consider $\{\varphi_k(j)\}_{k=1}^{4} = \{\sqrt{2}\sin(2\pi j), \sqrt{2}\cos(2\pi j), \sqrt{2}\sin(4\pi j), \sqrt{2}\cos(4\pi j)\}$.

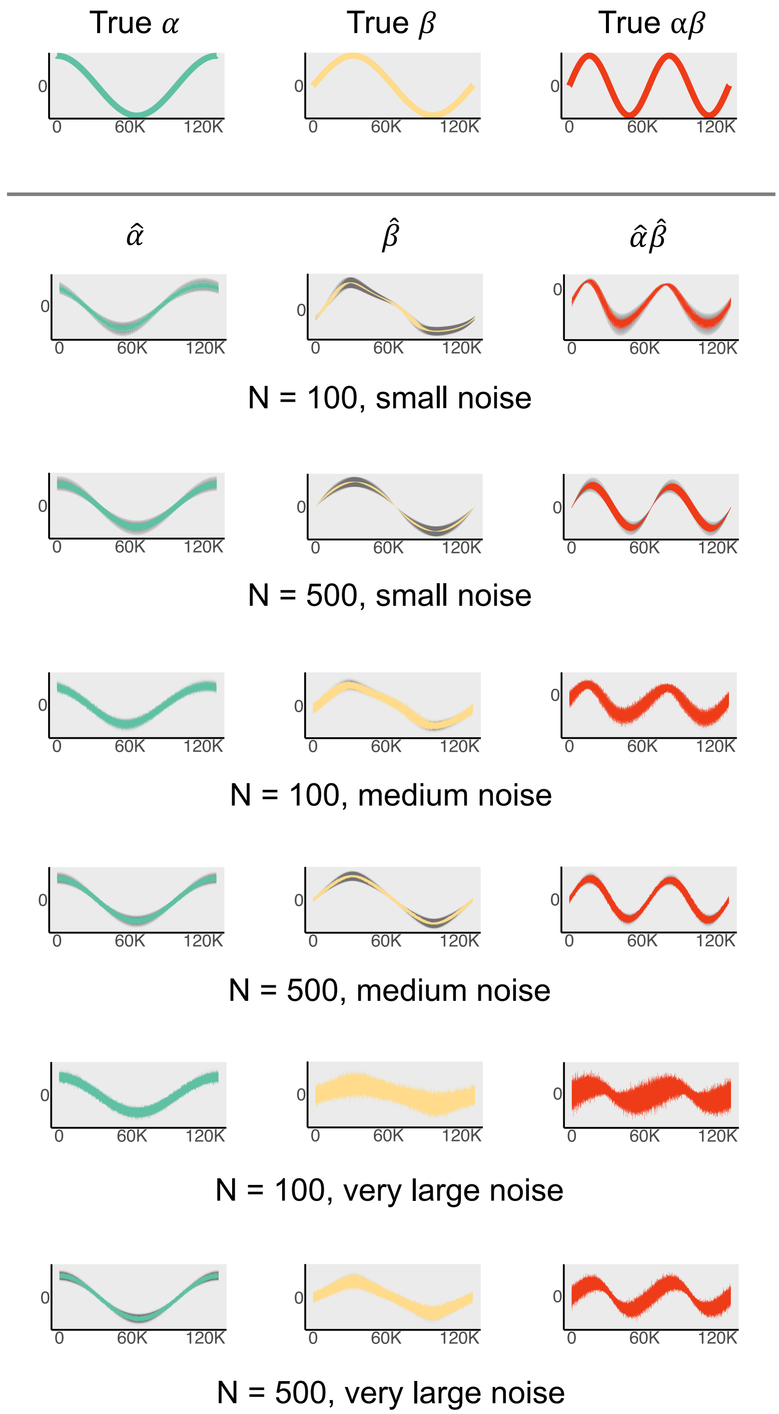


**Figure S1.** Simulation studies using orthogonal basis functions at different combinations of sample size and noise level.

#### 1.2 Non-orthogonal basis functions

Consider $\{\varphi_k(j)\}_{k=1}^{4} = \{1, \sqrt{3}(2j-1), \sqrt{5}(6j^2-6j+1), \sqrt{7}(20j^3-30j^2+12j-1)\}$.

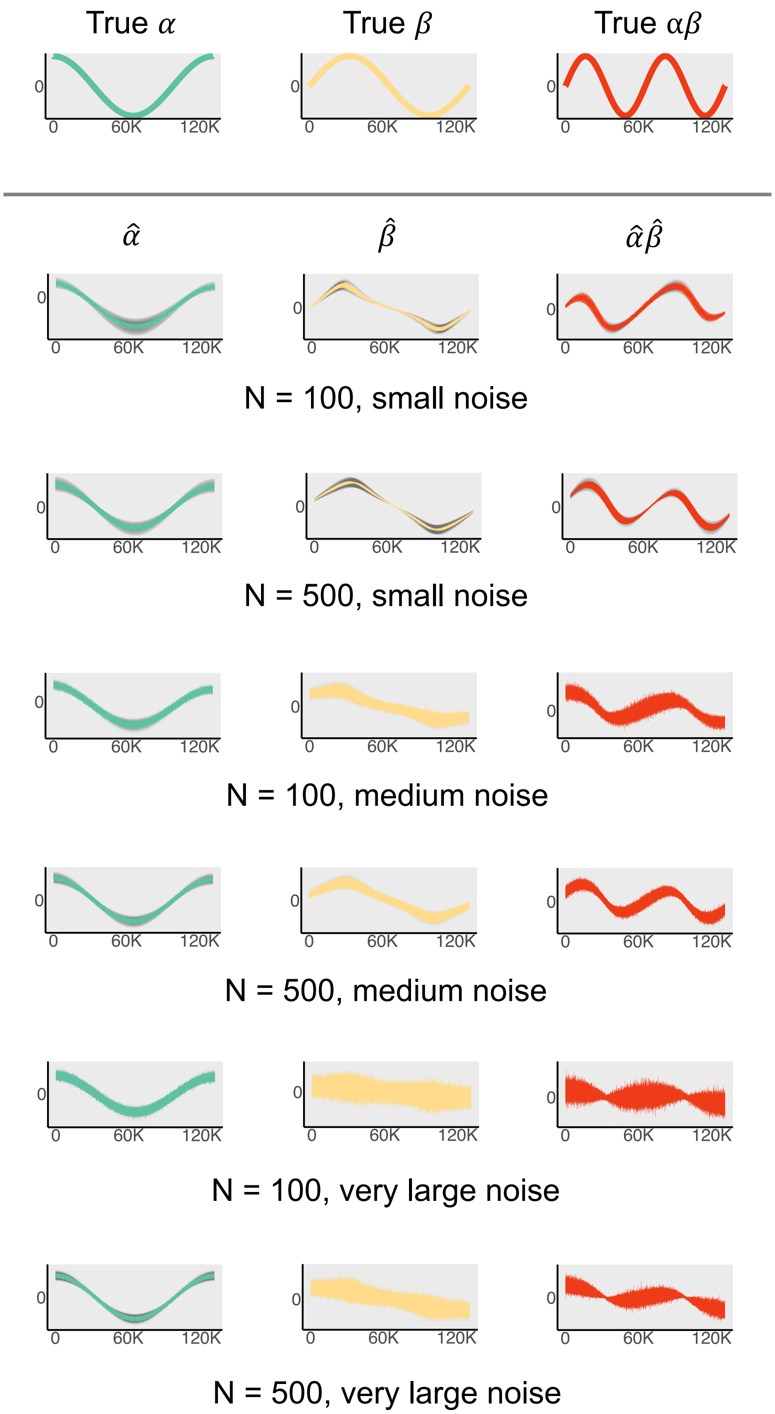


**Figure S2.** Simulation studies using nonorthogonal basis functions at different combinations of sample size and noise level.

### 2. Theoretical Properties

In the following, we demonstrate that the direct and indirect effects of a multivariate causal mediation model with binary outcomes can be captured on the odds ratio scale.

**Definitions.** Let $X$ denote an independent variable for a given subject (*e.g.*, prodromal symptoms), and $Y$ an outcome variable (*e.g.*, having psychosis or healthy). Suppose there are multiple mediators $\boldsymbol{M} = (M(1), M(2), \ldots, M(V))$. In the rs-fMRI context, the mediators are $V$ dependent activations over the $V$ voxels. Here we assume for simplicity that each subject is scanned under one condition. We use upper and lower cases to distinguish a random variable and its realization (real data), respectively. We use bold letter to specify a vector.

Using potential outcomes notation [1–3], let $Y(x) = Y(X = x)$, $\boldsymbol{M}(x) = \boldsymbol{M}(X = x)$, and $Y(x, \boldsymbol{m}) = Y(X = x, \boldsymbol{M} = \boldsymbol{m})$. In the following, we extend the direct and indirect effects defined on the odds-ratio for a single mediator [4] to incorporate multiple (and high-dimensional) mediators.

The total effect (TEs) conditional on a covariate $\boldsymbol{Z} = \boldsymbol{z}$ on the odds ratio (OR) is defined as

$$TE^{OR}_{x,x^*}|\boldsymbol{z} = \frac{P(Y(x)=1 \mid \boldsymbol{z})/\{1-P(Y(x)=1 \mid \boldsymbol{z})\}}{P(Y(x^*)=1 \mid \boldsymbol{z})/\{1-P(Y(x^*)=1 \mid \boldsymbol{z})\}} \qquad \text{(S.1)}$$

In words, it compares the odds of $Y$ being 1 if $X$ takes value $x$ and the odds of $Y$ being 1 had $X$ took value $x^*$, on the OR scale, in a given stratum where $\boldsymbol{Z} = \boldsymbol{z}$. In a psychosis study, this will capture the effect of symptom score changing from $x$ to $x^*$ on developing psychosis at the odds ratio scale, for a specific age, gender, and site group.

Next, the conditional controlled direct effects (CDEs) on the OR is defined as

$$CDE^{OR}_{x,x^*}|\boldsymbol{z} = \frac{P(Y(x,\boldsymbol{m})=1 \mid \boldsymbol{z})/\{1-P(Y(x,\boldsymbol{m})=1 \mid \boldsymbol{z})\}}{P(Y(x^*,\boldsymbol{m})=1 \mid \boldsymbol{z})/\{1-P(Y(x^*,\boldsymbol{m})=1 \mid \boldsymbol{z})\}} \qquad \text{(S.2)}$$

In words, it compares the odds of $Y$ being 1 if $X$ takes value $x$ and the high-dimensional mediator $\boldsymbol{M}$ is fixed (intervened to be) at $\boldsymbol{m}$, and the odds of Y being 1 had $X$ took value $x^*$ while $\boldsymbol{M}$ were fixed at $\boldsymbol{m}$, on the OR scale, in a given stratum where $\boldsymbol{Z} = \boldsymbol{z}$. In a psychosis study, this will capture the effect of symptom score changing from $x$ to $x^*$ on developing psychosis at the OR scale, with a brain region of interest fixed, via Transcranial magnetic stimulation (TMS), at $\boldsymbol{M} = \boldsymbol{m}$ for a specific age, gender, and site group.

Similarly, the conditional natural direct effects (NDEs) on the odds ratio is defined as

$$NDE^{OR}_{x,\underline{x}^*}|\boldsymbol{z} = \frac{P(Y(x,\boldsymbol{M}(x^*))=1 \mid \boldsymbol{z})/\{1-P(Y(x,\boldsymbol{M}(x^*))=1 \mid \boldsymbol{z})\}}{P(Y(x^*,\boldsymbol{M}(x^*))=1 \mid \boldsymbol{z})/\{1-P(Y(x^*,\boldsymbol{M}(x^*))=1 \mid \boldsymbol{z})\}} \qquad \text{(S.3)}$$

where the underlined $\underline{x}^*$ indicates that the mediator $\boldsymbol{M}$ is set at the level it would have been under $x^*$ despite the independent variable being $x$. Hence, the natural direct effect quantifies the effect of the independent variable (changing from $x$ to $x^*$) on the outcome on the OR scale, with mediator fixed at $\boldsymbol{M}(x^*)$ in a given stratum where $\boldsymbol{Z} = \boldsymbol{z}$.. In a psychosis study, this will capture the effect of symptom score changing from $x$ to $x^*$ on developing psychosis on the OR scale, with a brain region of interest fixed, via TMS, at the level had symptom score been $x^*$ (*i.e.* at $\boldsymbol{M}(x^*)$), for a specific age, gender, and site group.

Finally, the conditional natural indirect effects (NIEs) on the OR are defined as

$$NIE^{OR}_{\underline{x},x^*}|\boldsymbol{z} = \frac{P(Y(x,\boldsymbol{M}(x))=1 \mid \boldsymbol{z})/\{1-P(Y(x,\boldsymbol{M}(x))=1 \mid \boldsymbol{z})\}}{P(Y(x,\boldsymbol{M}(x^*))=1 \mid \boldsymbol{z})/\{1-P(Y(x,\boldsymbol{M}(x^*))=1 \mid \boldsymbol{z})\}} \qquad \text{(S.4)}$$

where the underlined $\underline{x}$ indicates that the independent variable $X$ is set at the level $x$ despite the mediator is set at level $\boldsymbol{M}(x^*)$ (namely, to what the mediator would have been had the independent variable been set to $x^*$). Hence, the natural indirect effect quantifies the effect of the mediator (changing from $\boldsymbol{M}(x)$ to $\boldsymbol{M}(x^*)$) on the outcome on the OR scale, in a given stratum where $\boldsymbol{Z} = \boldsymbol{z}$. In a psychosis study, this will capture the effect of brain measurement of a region of interest (changing from $\boldsymbol{M}(x)$ to $\boldsymbol{M}(x^*)$ while fixing symptom score at $x$) on developing psychosis on the OR scale, for a specific age, gender, and site group.

It follows, from Equations (S.1), (S.3), and (S.4), that the above total effect can be decomposed into a natural effect component and an indirect effect component on the OR scale:

$$TE^{OR}_{x,x^*}|\boldsymbol{z} = \{NIE^{OR}_{\underline{x},x^*} \times NDE^{OR}_{x,\underline{x}^*}\}\,|\boldsymbol{z}$$

The direct effect could also be defined as $Y\big(x,\boldsymbol{M}(x)\big) - Y\big(x^*,\boldsymbol{M}(x)\big)$. In general, this would lead to a different decomposition of the total effect; this, however, is not of further concern as the modification is straightforward.

**Identification.** To identify the direct and indirect effects using observed data, we require a few assumptions [5,6]. In the following, we assume $\boldsymbol{Z}$ contains all relevant confounding variables.

Assumption 1: $Y(x) \perp X \mid \boldsymbol{Z}$.

Assumption 2: $Y(x,\boldsymbol{M}(x)) \perp X \mid \boldsymbol{Z}$.

Assumption 3: $Y(x,\boldsymbol{M}(x)) \perp \boldsymbol{M} \mid \{X,\boldsymbol{Z}\}$.

Assumption 4: $\boldsymbol{M}(x) \perp X \mid \boldsymbol{Z}$.

Assumption 5: $Y(x,\boldsymbol{m}) \perp \boldsymbol{M}(x^*) \mid \boldsymbol{Z}$.

Conditioning on covariate $\boldsymbol{Z}$, these assumptions imply that, there is no confounding for the relationship between: (A1-2) independent variable $X$ and outcome $Y$; (A3) mediator $\boldsymbol{M}$ and outcome $Y$; (A4) independent variable $X$ and mediator $\boldsymbol{M}$; and (A5) no confounding for the relationship between mediator and outcome that is affected by the independent variable. Together, they are often referred to as sequential ignorability assumptions. See [7,8] for detailed discussion of these assumptions, and see [9] for a critical evaluation of these assumptions in the high-dimensional setting.

Under these assumption, [4,10] showed the average direct and indirect effects are identified from the regression function using the observed data for a single mediator and multiple mediators. Here, we show that their argument applies also to cases with high-dimensional mediators. In particular, under Assumption 1, the total effect conditional on a covariate $\boldsymbol{Z} = \boldsymbol{z}$ on the OR scale is identifiable:

$$TE^{OR}_{x,x^*}|\boldsymbol{z} = \frac{P(Y=1\,|x,\boldsymbol{z})/\{1-P(Y=1\,|x,\boldsymbol{z})\}}{P(Y=1\,|x^*,\boldsymbol{z})/\{1-P(Y=1\,|x^*,\boldsymbol{z})\}}.$$

Under Assumptions 2-3, the conditional direct effect on the OR scale is identifiable:

$$CDE^{OR}_{x,x^*}|\boldsymbol{z} = \frac{P(Y=1 \mid x, \boldsymbol{m}, \boldsymbol{z})/\{1-(Y=1 \mid x, \boldsymbol{m}, \boldsymbol{z})\}}{P(Y=1 \mid \ x^*, \boldsymbol{m}, \boldsymbol{z})/\{1-(Y=1 \mid \ x^*, \boldsymbol{m}, \boldsymbol{z})\}}.$$

Similarly, under Assumptions 2-5, the natural direct and indirect effects on the OR scales are identifiable.

Suppose Assumptions 2-5 hold, and the model in Equations (iv) in the main text is under a logit link function,

$$E(M(v)|X=x, \boldsymbol{Z}=\boldsymbol{z}) = \alpha_0 + \alpha(v)x + \boldsymbol{t}(v)\boldsymbol{z}, \quad v = 1, 2, \dots, V$$

$$logit(Y=1|X=x, \boldsymbol{M}=\boldsymbol{m}, \boldsymbol{Z}=\boldsymbol{z}) = \ \beta_0 + \gamma x + \sum\nolimits_{v=1}^{V} m(v)\beta(v) + \ \boldsymbol{\tau}^{\mathrm{T}}\boldsymbol{z}$$

The average controlled direct effect and natural indirect effect on the OR scale can be shown to be:

$$CDE^{OR}_{x,x^*}|\boldsymbol{z} = e^{\gamma(x-x^*)} \qquad \text{(S.5)}$$

$$NIE^{OR}_{\underline{x},x^*}|\boldsymbol{z} = e^{[\sum_{v=1}^{V} \alpha(v)\beta(v)](x-x^*)} \qquad \text{(S.6)}$$

Throughout, we assume that the independent variable does not interact with one or more of the mediators. When the independent variable interacts with one or more of the mediators, however, the bimodal framework considered in this paper is not appropriate for mediation analysis [11]. Instead, one may consider:

$$E(M(v)|X=x, \boldsymbol{Z}=\boldsymbol{z}) = \alpha_0 + \alpha(v)x + \tilde{\boldsymbol{t}}(v)\boldsymbol{z}, \quad v = 1, 2, \dots, V$$

$$logit(Y=1|X=x, \boldsymbol{M}=\boldsymbol{m}, \boldsymbol{Z}=\boldsymbol{z}) = \ \beta_0 + \gamma x + \sum\nolimits_{v=1}^{V} m(v)\beta(v) + \sum\nolimits_{v=1}^{V} x\, m(v)\, b(v) + \ \tilde{\boldsymbol{\tau}}^{\mathrm{T}}\boldsymbol{z}$$

Assume Assumptions 2-3 hold, we have

$$CDE^{OR}_{x,x^*}|\boldsymbol{z} = e^{[\gamma + \sum_{v=1}^{V} m(v)\, b(v)](x-x^*)}$$

$$NIE^{OR}_{\underline{x},x^*}|\boldsymbol{z} = e^{[\sum_{v=1}^{V} \alpha(v)\beta(v) \,+\, \sum_{v=1}^{V} \alpha(v)\, b(v)x](x-x^*)}$$

**Remarks.** When the counterfactuals are well defined and the Assumptions 1-5 hold, the right-hand side of Equations S.5 and S.6 identify causal mediation effects. When one or more of the assumptions in Assumptions 1-5 fail to hold, or if the counterfactuals are not well defined, the right-hand side of Equations S.5 and S.6 may still be used in exploratory analysis to help identify potential mediators. For example, they could identify linear combinations of voxels that correspond to specific brain functions, suggesting mediation through correlates of those brain functions. In the article, for simplicity, we use “direct effect” and “indirect effect” to refer to the right-hand sides of Equations S.5 and S.6, respectively; we are agnostic throughout as to whether these expressions can be interpreted causally or should be taken as exploratory.

Similarly, we use “mediator” agnostically to refer to variables that temporally follow the independent variable and precede outcome and potentially may lie on a causal pathway between them. We also note that the model only considers the case where the entire vector $\boldsymbol{M}$ is subject to a single level of intervention ($x$). More generally, it would be possible to define potential mediators when some elements of $\boldsymbol{M}$ are subject to $x$ and other elements of $\boldsymbol{M}$ are subject to $x^*$, which could support various decompositions of direct/indirect effects due to pathways acting through different $\boldsymbol{M}$. In the fMRI setting described in the article we anticipate that the same elements of $\boldsymbol{M}$ (brain regions) are subject to both $x$ and $x^*$. For example, minor prodromal symptoms and severe prodromal symptoms are associated with activation in the same brain regions, but with different intensities.

### 3. A Comparison between Different Mediation Analysis Approaches

To complement the presentations made in this study, in the following we provide a brief comparison between common mediation analysis. To avoid further confusion, we suppose that all covariates have been controlled.

There are, broadly, two approaches to conduct mediation analyses: the *difference* approach and the *product* approach [12,13]. Conceptually, both approaches employ a series of regression models to dissect the pathways between an independent variable, a mediator, and an outcome.

The difference approach determines that a variable $M$ is a mediator as follows. Consider two regression models, one consisting of regressing an independent variable $X$ on the outcome $Y$ (call the estimated effect from the independent variable to the outcome as $\gamma$), and the other consisting of regressing both the independent variable $X$ and the potential mediator $M$ on the outcome $Y$ (call the estimated effect from the independent variable to the outcome under this setting as $\gamma'$). If the effect of the independent variable $X$ on the outcome $Y$ when the mediator $M$ is not included in the regression model is considerably greater than the effect of the independent variable $X$ on the outcome $Y$ when the potential mediator $M$ is included, it suggests that there is a significant mediation effect (that alleviates the direct effect of the independent variable on the outcome), and thus the mediation effect can be estimated by taking the difference between $\gamma$ and $\gamma'$.

The product approach also considers two regression models, one consisting of regressing the independent variable $X$ on the potential mediator $M$ (call the effect of the independent variable on the potential mediator as $\alpha$), and the other consists of regressing both the independent variable $X$ and the potential mediator $M$ on the outcome $Y$ (call the effect of the potential mediator on the outcome while controlling the independent variable as $\beta$). The product approach determines that $M$ is a mediator if both $\alpha$ and $\beta$ effects are significantly different from zero. Then, it suggests that there exists a pathway from the independent variable $X$ to the outcome $Y$, connected (mediated) by $M$, and the mediation effect can be estimated by taking the product of $\alpha$ and $\beta$.

When the outcome is continuous, the estimated mediation effects from the difference and the product approaches are the same; when the outcomes are binary (such as being healthy or afflicted with a certain type of disease), and when the (disease) rate is rare ($\leq 10\ \%$), the product approach and difference approach approximately coincide [4]. When the outcomes are binary, but the (disease) rate is common, the product approach and the difference approach diverge. The difference approach becomes particularly challenging because the coefficient of the independent variable using this method may not change when there is mediation (see [12] and see Chapter 2 of [14] for a discussion).

Since the product approach is suitable for both rare and common cases (the dataset used in this study contains rare outcomes), we adopt this approach in this article. For the sake of completeness, we provide a brief technical summary of both approaches below.

**The product approach.** The product mediation analysis framework involves two steps [15–19]. Consider the setting in **Figure 1** (a) of the main text. The first step examines if the independent variable $X$ is associated with the mediator $M$. This is done using a conditional regression model:

$$E(M|x) = \theta_0 + \alpha x \quad \text{(S.7)}$$

where $E$ refers to the expectation operation, and $\theta_0$ and $\alpha$ are coefficients for the intercept, and the independent variable $x$. In simpler terms, if the coefficient $\alpha$ in (S.7) is significantly different from zero, it suggests that there is potentially an effect from the independent variable to the mediator. In other words, the red arrow below *alpha* in **Figure 1** (a) exists.

The second step checks if the mediator $M$ is associated with the outcome $Y$, after controlling for the independent variable $X$. This is done using another conditional regression model:

$$E(Y|x,m) = \theta_0' + \gamma x + \beta m \quad \text{(S.8)}$$

where $\theta_0'$, $\gamma$, and $\beta$ are coefficients for the intercept, the independent variable $x$, and mediator $m$. In simpler terms, if the coefficient $\beta$ in (S.8) is significantly different from zero, it suggests that there is potentially an effect from the mediator to the outcome. In other words, the red arrow below *beta* in **Figure 1** (a) exists.

Taken together, when both $\alpha$ and $\beta$ are significantly different from zero, it means that (I) a change of $X$ affects $M$, and (II) a change in $M$ further affects $Y$; therefore $M$ is significantly mediating the relationship between $X$ and $Y$. In simpler terms, the mediation effect exists when the product $\alpha\beta$, which quantifies the mediation effect in this setting, is non-zero. In other words, both red arrows below *alpha* and *beta* in **Figure 1** (a) exist, thereby connecting $X$ and $Y$ via a pathway that passes through (mediated by) $M$. The remaining coefficient $\gamma'$ in (S.8) quantifies the direct effect of the independent variable $X$ on outcome $Y$ (that is not mediated by $M$) (see **Figure 1** (a)).

**The difference approach.** The difference approach has two modeling steps. The first step is a conditional regression model, which investigates if variability in the independent variable $X$ is responsible for variability in outcome $Y$.

$$E(Y|x,\mathbf{z}) = \theta_0 + \gamma x \quad \text{(S.9)}$$

where $E$ stands for expectation, $\theta_0$ and $\gamma$ are coefficients for the intercept and the independent variable $x$.

The second step is a conditional regression model, which checks if variability in the independent variable $X$ and mediator $M$ are both responsible for variability in outcome $Y$.

$$E(Y|x,m) = \theta_0' + \gamma' x + \beta m \quad \text{(S.10)}$$

where $\theta_0'$, $\gamma'$, and $\beta$ are coefficients for the intercept, the independent variable $x$, and mediator $m$. Note the coefficients are denoted with a prime (*e.g.* $\gamma'$ in (S.10)) to highlight that they are not necessarily the same as their counterparts (*i.e.* $\gamma$ in (S.9)).

If the coefficient of the independent variable on the outcome when a mediator is included, or $\gamma'$, is considerably smaller than the coefficient of the independent variable on the outcome when the mediator is not included, or $\gamma$, then it suggests that the mediator is at least partially responsible for explaining the variance (information) of the outcome $Y$. To put more concretely, this indicates that including the mediator $m$, the independent variable $x$ explains a less amount of variance (information) of $Y$. The difference of $\gamma - \gamma'$ thus quantifies the mediation effect in this setting.

### 4. A Brief Procedure for Bootstrap Experiments Used in This Study

Denote $(\boldsymbol{x}_1^*, \boldsymbol{y}_1^*, \boldsymbol{M}_1^*)$, $(\boldsymbol{x}_2^*, \boldsymbol{y}_2^*, \boldsymbol{M}_2^*), \dots, (\boldsymbol{x}_B^*, \boldsymbol{y}_B^*, \boldsymbol{M}_B^*)$ as *B* sets of bootstrap data, where the independent variable, the outcome variable, and the mediator are jointly sampled from ($\boldsymbol{x}$, $\boldsymbol{y}$, $\boldsymbol{M}$). The $j^{th}$ bootstrap sample has the same dimension as ($\boldsymbol{x}$, $\boldsymbol{y}$, $\boldsymbol{M}$); namely $\boldsymbol{x}_j^*$ and $\boldsymbol{y}_j^*$ each contains $N$ bootstrapped observations and $\boldsymbol{M}_j^*$ is $N \times V$. Let $NIE_1^*(j), NIE_2^*(j), \dots, NIE_B^*(j)$ be the estimated mediation effect from each bootstrap experiment for the $j^{th}$ voxel. Specifically, $NIE_b^*(j)$ is the estimated indirect mediation effect corresponding to the $j^{th}$ voxel using data $(\boldsymbol{x}_b^*, \boldsymbol{y}_b^*, \boldsymbol{M}_b^*)$.

The bootstrap *t*-statistic for the mediation effect corresponding to the $j^{th}$ voxel is defined as

$$t_j^* = \frac{\overline{NIE}^*(j)}{SE_{NIE^*(j)}}$$

where $\overline{NIE}^*(j) = \frac{\sum_{i=1}^{B} NIE_i(j)}{B-1}$ is the mean indirect mediation effect for the $j^{th}$ voxel, averaged across *B* bootstrap experiments, and $SE_{NIE^*(j)} = \frac{\sum_{i=1}^{B}(NIE_i^*(j) - \overline{NIE}^*(j))^2}{\sqrt{B}}$ is the standard error.

### References


1. Rubin, D. B. Estimating causal effects of treatments in randomized and nonrandomized studies. *J. Educ. Psychol.* **66**, 688–701 (1974).
2. Albert, J. M. Mediation analysis via potential outcomes models. *Stat. Med.* **27**, 1282–1304 (2008).
3. Imai, K., Jo, B. & Stuart, E. A. Commentary: Using potential outcomes to understand causal mediation analysis. *Multivariate Behav. Res.* **46**, 861–873 (2011).
4. VanderWeele, T. J. & Vansteelandt, S. Odds ratios for mediation analysis for a dichotomous outcome. *American Journal of Epidemiology* **172**, 1339–1348 (2010).
5. Pearl, J. Direct and indirect effects. In *Proc. seventeenth Conf. Uncertain. Artif. Intell.*, edited by J. Breese & D. Koller, pp. 411–420. Morgan Kaufmann Publishers, San Francisco, CA, USA (2001).
6. Vanderweele, T. J. & Vansteelandt, S. Conceptual issues concerning mediation, interventions and composition. *Stat. Interface* **2**, 457–468 (2009).
7. Robins, J. M. & Richardson, T. S. Alternative graphical causal models and the identification of direct effects. *In Causality and Psychopathology: Finding the Determinants of Disorders and Their Cures, edited by P.E. Shrout, K.M. Keyes & K. Ornstein. Oxford University Press, Oxford, UK* (2010).
8. Pearl, J. Interpretation and identification of causal mediation. *Psychol. Methods* **19**, 459–481 (2014).
9. Huang, Y. T. & Pan, W. C. Hypothesis test of mediation effect in causal mediation model with high-dimensional continuous mediators. *Biometrics* **72**, 402–413 (2016).
10. VanderWeele, T. J. & Vansteelandt, S. Mediation Analysis with Multiple Mediators. *Epidemiol. Method.* **2**, 95–115 (2014).
11. Ogburn, E. L. Commentary of 'Mediation Analysis Without Sequential Ignorability: Using Baseline Covariates Interacted with Random Assignment as Instrumental Variables' by Dylan Small. *J. Stat. Res.* **46**, 105–111 (2012).
12. VanderWeele, T. J. Mediation Analysis: A Practitioner's Guide. *Annu. Rev. Public Health* **37**, 17–32 (2016).
13. Mackinnon, D. P. & Dwyer, J. H. Estimating Mediated Effects in Prevention Studies. *Eval. Rev.* **17**, 144–158 (1993).
14. VanderWeele, T. J. Explanation in causal inference: Developments in mediation and interaction. *Int. J. Epidemiol.* **45**, 1904–1908 (2016).
15. Hyman, H. H. *Survey Design and Analysis: Principles, Cases and Procedures*. (Free Press, 1955).
16. Alwin, D. F. & Hauser, R. M. The Decomposition of Effects in Path Analysis. *Am. Sociol. Rev.* **40**, 37 (1975).
17. Judd, C. M. & Kenny, D. A. Process analysis: Estimating Mediation in Treatment Evaluations. *Eval. Rev.* **5**, 602–619 (1981).
18. Sobel, M. E. Asymptotic confidence intervals for indirect effects in structural equation models. *Sociol. Methodol.* **13**, 290–312 (1982).
19. Baron, R. M. & Kenny, D. a. The moderator-mediator variable distinction in social psychological research: conceptual, strategic, and statistical considerations. *J. Pers. Soc. Psychol.* **51**, 1173–1182 (1986).
